# CYP2C19*2, CYP2C19*17 and ITGB3 (PlA1/A2) polymorphisms and resulting therapeutic alterations of clopidogrel and aspirin: A clinical screening among CVD patients who undergone PCI

**DOI:** 10.1101/2020.10.04.325258

**Authors:** Tasnova Tasnim Nova, Ishrat Jahan, N.A.M. Momenuzzaman, Sikder Nahidul Islam Rabbi, Thomas M.C. Binder, Mohammad Safiqul Islam, Md. Rabiul Islam, Abul Hasnat, Zabun Nahar

## Abstract

**Introduction:** Clopidogrel and aspirin are at the base of treatment in conditions like arterial thrombosis and after patients have undergone percutaneous coronary intervention. But frequently found CYP2C19*2 and CYP2C19*17 polymorphisms and some variants of the ITGB3 gene cause alteration in the therapeutic effectiveness of this drug.

**Methods:** One thousand cardiovascular patients were recruited for each drug under study. Their blood was collected to analyze the genotype using PCR-RFLP and T-ARMS-PCR method for clopidogrel and aspirin respectively. The PCR products for clopidogrel were screened with agarose gel electrophoresis and then digested with SmaIfor CYP2C19*2 and Nsil-HF for CYP2C19*17. The digested products of clopidogrel and the ARMS-PCR product of aspirin were run on 2% AGE to analyze the polymorphisms.

**Results:** In our outcome, the percentage of hetero and mutant homozygous people in CYP2C19*2 polymorphism (loss-of-function allele) was 64.1% and for CYP2C19*17 (gain-of-function allele) was 22.3%. For ITGB3 polymorphism, it was found that 84.1% of them belonged to the homozygous group while 15.6% was heterozygous and only 0.3% were mutant homozygous patients.

**Conclusion:** Our study findings were quite compatible with the results of some other studies in other ethnic groups. This phenomenon suggested for modification of dose or application of alternative generics in patients who are under the risk of therapeutic failure or toxicity produced by these drugs.

## 1. Introduction

Cardiovascular diseases (CVD) are of immense clinical importance and pharmacogenomics implications [1-3]. In 2013, CVD was the most common underlying cause of death in the world, accounting for an estimated 17.3 million (95% clearance interval, 16.5-18.1 million) of 54 million total deaths, or 31.5% (95% clearance interval, 30.3%-32.9%) of worldwide deaths [4]. Coronary thrombosis accounts for one of the prime causes of mortality in CVD. Patients suffering from acute coronary syndromes (ACSs) undergoing percutaneous coronary intervention (PCI) require adequate platelet inhibition to prevent recurrent ischemic events [8]. Oral antiplatelet drugs, clopidogrel, and aspirin are the foundation and the most widely used mode of treatment in conditions like arterial thrombosis [5]. These medications are widely used for the management of ACS and stent thrombosis [6]. As regards for patients undergoing carotid endarterectomy, perioperative antithrombotic therapy can be composed of aspirin alone, while high-risk patients and drug-eluting plaques are managed with aspirin plus clopidogrel, ticagrelor, or prasugrel for a recommended duration [7, 8].

Clopidogrel is a second-generation oral thienopyridine which is a prodrug and is converted into its active metabolite by hepatic cytochrome (CYP) P450 [6]. It has been demonstrated by several pharmacogenomic studies that CYP2C19 variant alleles affect the pharmacokinetics and pharmacodynamics of clopidogrel and influence the inter-individual heterogeneity in clopidogrel response [6-12]. Variant alleles accounting for lessened function or loss of function have been shown to encompass CYP2C19*2 to CYP2C19*8 [6]. The CYP2C19*2 gene variant is a G681A mutation in exon 5 that encodes for a hidden link variant, while CYP2C19*3 a G636A mutation in exon 4 results in a premature stop codon [7]. The CYP2C19*17 allelic variant, a C806T mutation in exon 5, is responsible for increased catalytic activity. Carriage of CYP2C19*17 allele results in a higher platelet inhibition by clopidogrel owing to an augmented form of the active metabolite which could result in an increased risk of bleeding [11]. The gain-of-function CYP2C19*17 T allele always occurs on a haplotype that also shelters the wild-type “CYP2C19 *2 G”, so it can be assumed that the spotted effect of the gain-of-function *17 alleles may be due, in part, to the absence of the loss-of-function CYP2C19 *2 A allele [12]. Hitherto, studies have examined whether clopidogrel treatment is affected by CYP2C19 variant alleles in patients with acute coronary syndrome [13,14], myocardial infarction (MI) [15], coronary artery disease [16, 17] and patients undergoing PCI [18-22]. According to published data, patients who are homozygous for nonfunctional alleles of the CYP2C19 gene are termed “CYP2C19 poor metabolizers”. Nearly 2% of white and 4% of black patients are poor metabolizers; the prevalence of poor metabolizers is higher in Asian patients (14% of Chinese) [23-26].

Aspirin is another antiplatelet agent widely used for the prevention of ischemic cardiovascular and cerebrovascular events. The long-term dispensation of aspirin to high-risk CVD patients results in a 25% reduction in atherothrombotic stroke, MI, and death. Along with this it also caused a decline in the rate of bypass surgery, pulmonary embolism events, arterial thromboembolic events, and deep vein thromboses [27]. Aspirin resistance occasions the growth of thrombotic plaques during treatment with the drug and the ITGB3 polymorphism of the platelet receptor glycoprotein IIIa is the most frequently studied genetic variant and an important factor contributing towards its biological resistance [28]. This polymorphic ITGB3 gene at codon 33 of exon 2 (T1565C, rs5918) corresponds to the polymorphism of amino acid residues (leucine/proline) at position 33 of the polypeptide chain modulates platelet function while the PlA2 allele is associated with amplified platelet reactivity [29]. Ferguson *et. al*., Pamukcu*et. al*. and Angiolillo *et. al*. also concluded that there is a genetic association with the hyporesponsiveness to antiplatelet therapy [30-32].

The combined therapy of aspirin and clopidogrel deter small particles in the blood from adhering together and forming clots, so bleeding incidents were tracked to see if paired therapy was safe. A study of the American Heart Association demonstrated the efficacy and safety profile of the combined clopidogrel-aspirin therapy which imparts better protection against subsequent stroke [33]. The employment of clopidogrel as a part of the dual antiplatelet scheme represents a standard of care in clinical practice [3-4,9]. Around 40 million patients worldwide are using the antiplatelet effect of clopidogrel to treat or prevent atherothrombotic events and after percutaneous coronary revascularization [10]. However, the linkage between the occurrence of post-PCI ischemic events and clopidogrel therapy demonstrated a widely uneven pharmacodynamic response whereby approximately 1 in 3 patients has high on-treatment platelet reactivity (HTPR) [3]. Regardless of their proven clinical effect, a considerable number of patients do not have a satisfactory response to clopidogrel [9]. About 10 to 15% of patients receiving this medication have periodic atherothrombotic events [7,10].

On knowing the predicament about clopidogrel, the U.S. Food and Drug Administration imparted a warning suggestive of diminished effectiveness of standard doses of the drug in individuals with 2 reduced-functional CYP2C19 alleles and of genetic tests useful to set apart patients who are at risk and to think over alternative treatment policies [34]. Substantial clinical trials that evaluated higher loading and maintenance dose of clopidogrel in ACS/PCI patients have resolved that adjusting the dose does not downgrade the incidence of death [35]. Prasugrel and ticagrelor, on the other hand, were superior to clopidogrel in a large-scale randomized trial of ACS patients with planned PCI which showed 42% and 26 % reduction in-stent thrombosis, respectively [36]. However, these may not replace clopidogrel for all kinds of patients due to contraindication in some patients [36], insufficient long-term toxicity data [37] and also lack of statistically significant data [38].

As of April 2013, clinical pharmacogenetics implementation consortium (CPIC) guideline recommended the application of an updated therapeutic approach based on a prospective randomized controlled trial on CYP2C19 genotype-directed antiplatelet therapy especially on patients with ACS undergoing PCI [39]. Considering all these aspects, we had therefore undertaken this study to find out the prevalence of these SNPs in CVD patients undergone PCI and their relationship with in vivo inactivation of clopidogrel. We plan to further extend the study to analyze for the metabolites produced in patients with variant genetic make-up and suggest an alternative molecule or mode of treatment among the Bangladeshi population for the very first time.

## 2. Methods

### 2.1 Subject selection

A screening study was conducted on 1000 cardiovascular patients for clopidogrel and 1000 patients for aspirin. Patients were recruited from the Cardiology Department of United Hospital who have undergone percutaneous coronary intervention between March 2017 and November 2018. The study was conducted following the Helsinki Declaration and its further amendments [38,39]. Written ethical permission from the ethical review committee of the respective hospital was taken to approve the protocol. Before the study, all the patients had given written consent in a prescribed form. This genetic study was conducted in the pharmacogenetics laboratory of Labaid Limited (Diagnostic), Dhaka, Bangladesh.

### 2.2 DNA extraction

DNA was extracted (using the Sacace blood DNA extraction kit, based on silica column technology) from whole blood collected in EDTA tube. The protocol was prefixed within the DNA extraction kit package [40]. To assess the purity of extracted DNA, the ratio of absorbances was measured for A260/280 by Eppendorf bio photometer with a single used cell.

### 2.3 Polymerase chain reaction

Total PCR reaction volume for clopidogrel was 10 µL, containing 5 µL master mix, 0.15 µL forward primer (FP) and 0.15 µL reverse primer (RP), 2.7 µL NFW (nuclease-free water) and 2µl DNA. PCR was done in the T100 thermocycler of Bio-rad.

Similarly for aspirin, the PCR reaction volume was also 10 µL whereas 5 µL master mix, 0.15 µL of each primer (forward outer+reverse outer; forward inner+reverse inner),2.4 µL NFW and 2µl DNA. Using the tetra-primer amplification refractory mutation system (T-ARMS-PCR), this PCR mixture was also run in T100 thermocycler Bio-rad.

### 2.4 The material used in the reaction

Promega G2 green Taq 2x master mix (Promega, USA) = 5 µL for each reaction. For clopidogrel, Oligos are from Bio-basic Canada, diluted to 10 times for working conditions and used in 10pmole/ µL concentration. 0.15 µL for each reaction (FP+RP). For aspirin, oligos are from Bio-basic Canada, diluted to 10 times for working conditions and used in 10pmole/ µL concentration. 0.15 µL for each reaction (Fo+Ro; Fi+Ri).

### 2.5 Genotyping

Genotyping of clopidogrel was done with PCR-RFLP method [41, 42] and screened with agarose gel electrophoresis (AGE) (1.5%) and later on digested with SmaI (For CYP2C19*2 polymorphism, incubated on 25 ^°^C for overnight) and Nsil-HF (For CYP2C19*17 polymorphism, incubated at 37 ^°^C for an hour). The digested product was run on 2% AGE to detect the mentioned polymorphisms. On the other hand, the genotyping of aspirin was done with T-ARMS-PCR for missense rs5918 (PlA1/A1) polymorphism of the ITGB3 gene [43]. Then the PCR product was analyzed with 2% AGE.

## 3. Results

One thousand patients were genotyped for each polymorphism in CYP2C19 and ITGB3 genes to investigate the response of clopidogrel and Aspirin, respectively, the most commonly prescribed anticoagulants in CVD patients in Bangladesh. In case of clopidogrel, digestion of CYP2C19*2 with SmaI resulted in products of 170bp and 120bp (homozygous, GG); 300bp, 170bp and 120bp (heterozygous, GA) and 300bp (mutant, AA) (Fig. 1) while CYP2C19*17 with NsiI resulted in 116bp (homozygous, CC); 143bp, 116 bp and 27bp (heterozygous, CT) and 143bp (mutant, TT) (Fig.2).

**Fig. 1.**
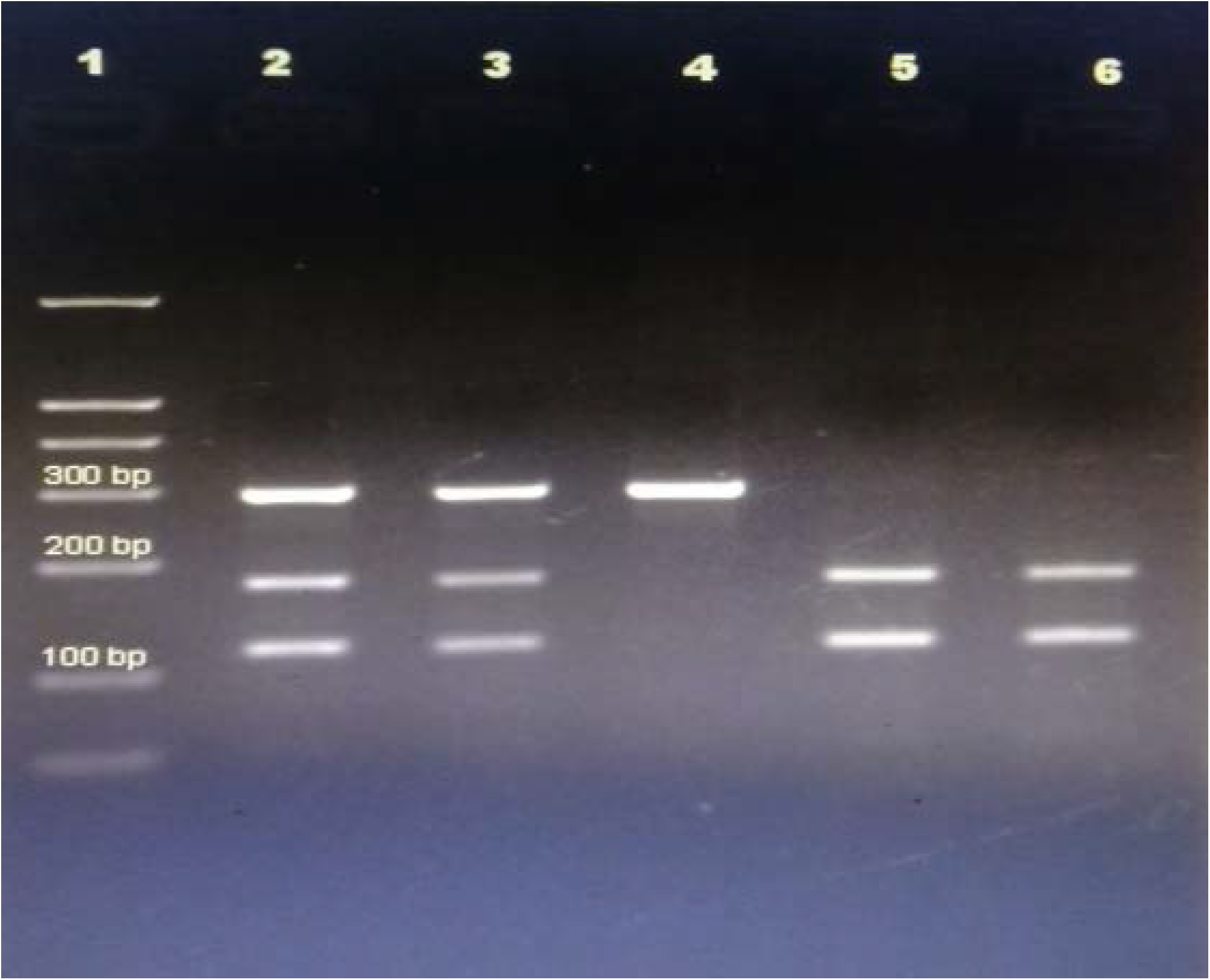
PCR-RFLP based genotyping of CYP2C19*2 (Lane 1 DNA ladder; lane 2,3 Heterozygous; lane 4 mutant and lane 5, 6 homozygous)

**Fig. 2.**
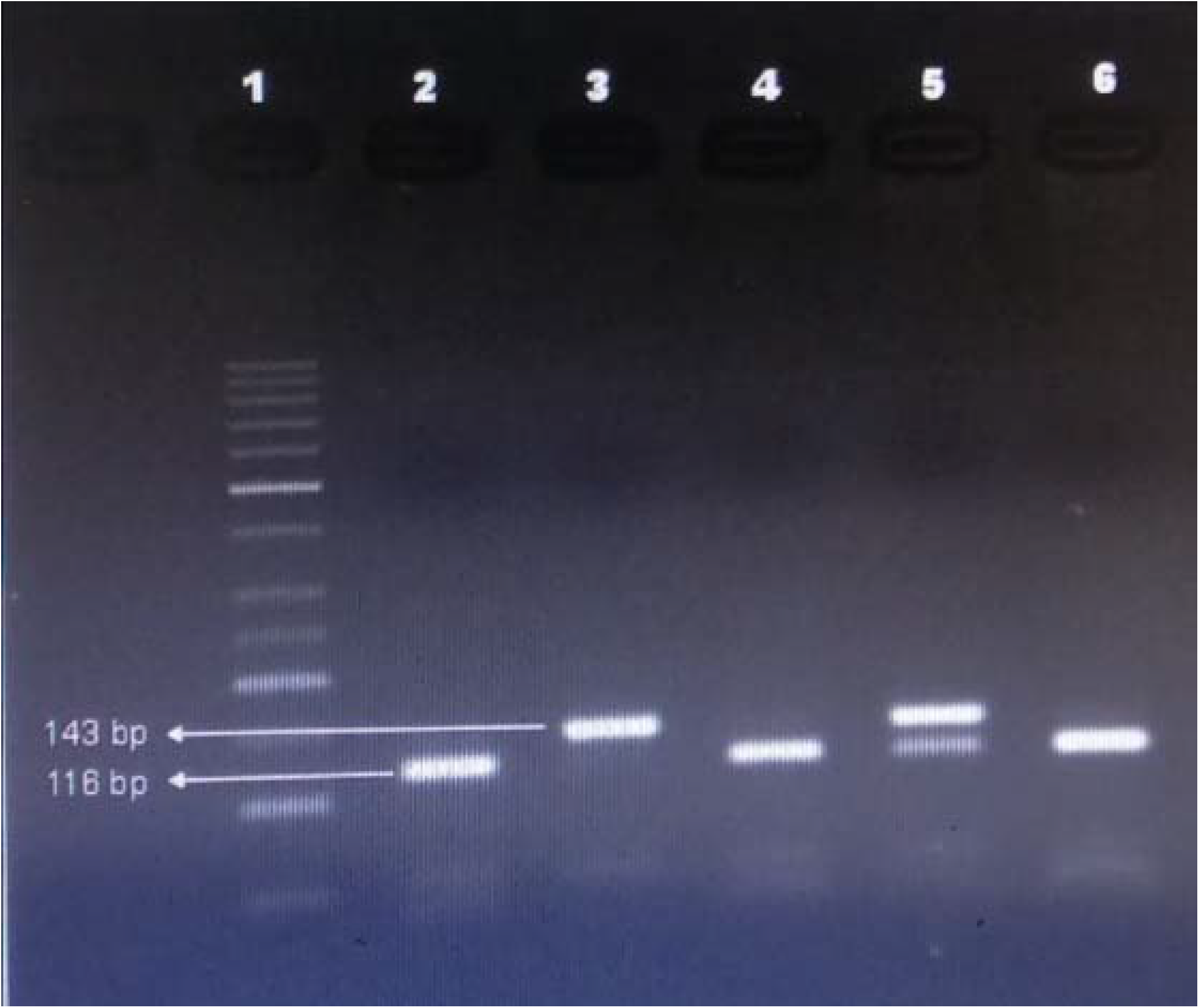
PCR-RFLP based genotyping of CYP2C19*17 (Lane 1 DNA Ladder; lane 2, 4, 6 homozygous; lane 3 mutant and lane 5 heterozygous)

In our findings, it was noticed that the percentage of hetero and mutant homozygous carriers PCI patients in CYP2C19*2 polymorphism is 64.1% which is a large number. But the percentage for the same variants found in CYP2C19*17 is quite less (22.7%). Still, both can be major issues while prescribing these two drugs in CVD patients who have undergone PCI. The percentage of homozygote in CYP2C19*2 is low (35.9%) and that is for CYP2C19*17 is 77.3%. (Table 1).

**Table 1.** Percentages of homozygous, heterozygous and mutant homozygous people in response to clopidogrel

| CYP2C19 | *2 |  | *17 |  |
| --- | --- | --- | --- | --- |
|  | *n* =1000 | % | *n* =1000 | % |
| Homozygous | 359 | 35.9 | 773 | 77.3 |
| Heterozygous | 472 | 47.2 | 204 | 20.4 |
| Mutant homozygous | 169 | 16.9 | 23 | 2.3 |
| Hetero + mutant | 641 | 64.1 | 227 | 22.7 |

Abbreviations: *CYP* Cytochrome, *n* Number.

On the other hand, in aspirin, 2 bands of homozygotes (PlA1/A1) resulting in 424bp and 285 bp; 3 bands in heterozygotes (PIA1/A2) were in 424, 285 and 180 bp and 2 bands of mutant (PIA2/A2) found in 424 and 180 bp after T-ARMS-PCR based genotyping of ITGB3 gene (Fig. 3).

**Fig. 3.**
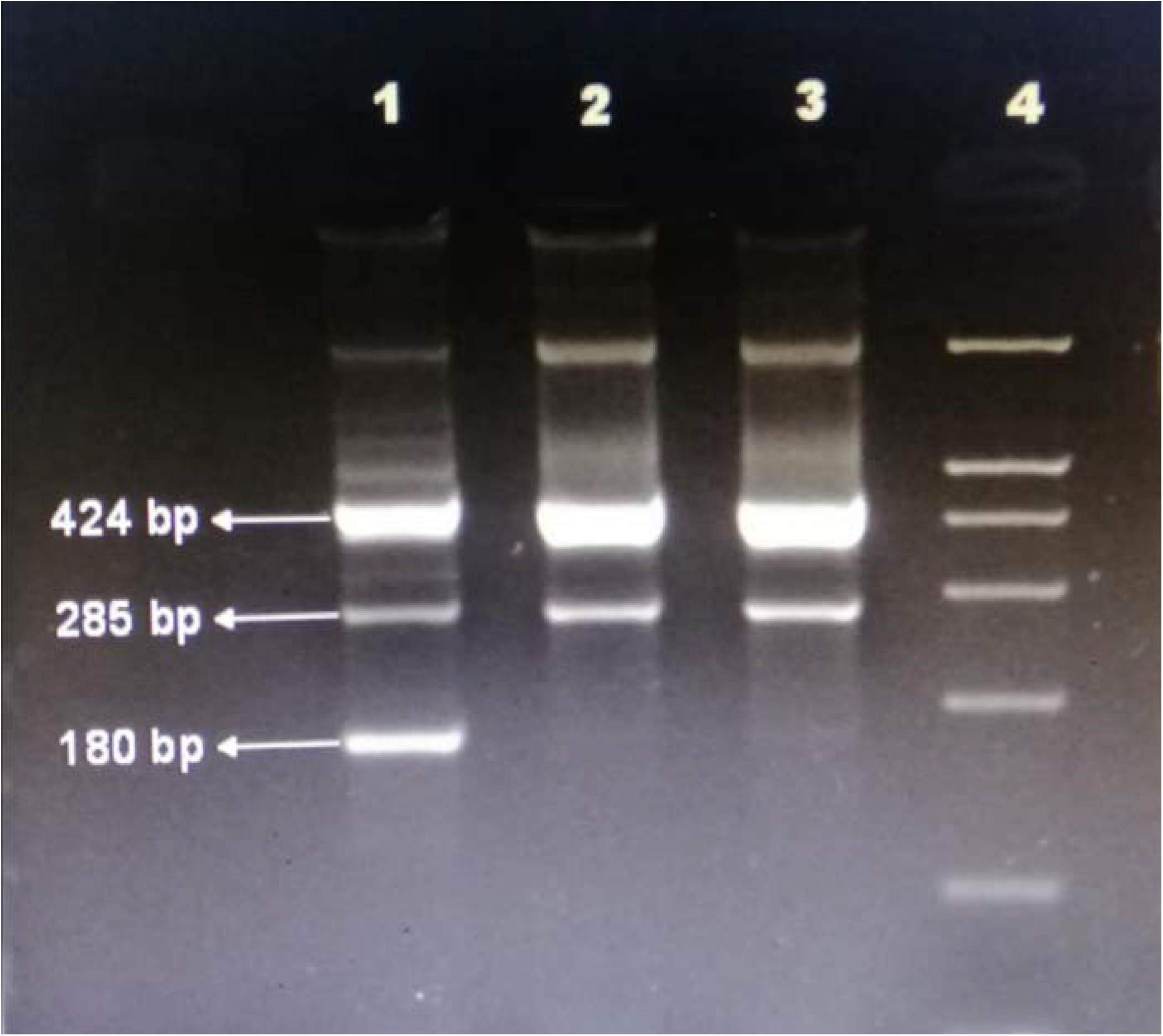
T-ARMS- PCR based genotyping of the ITGB3 gene (Lane 1 heterozygous; lane 2,3 normal and lane 4 DNA ladder)

The situation is not as devastating in the case of aspirin as in clopidogrel. As after examining 1000 patients for PlA1/A2 polymorphism of the ITGB3 gene, it was found that 84.1% of them belong to the homozygous (PlA1/A1) group leaving 15.6% heterozygous (PlA1/A2) and only 0.3% mutant (PlA2/A2) patients. The cumulative percentage for heterozygous and mutant homozygous patients was15.9 which is not an upsetting figure yet (Table 2).

**Table 2.** Percentages of homozygous, heterozygous and mutant homozygous people in response to aspirin

| ITGB3 | *n* =1000 | % |
| --- | --- | --- |
| Homozygous (PlA1/A1) | 841 | 84.1 |
| Heterozygous (PlA1/A2) | 156 | 15.6 |
| Mutant (PlA2/A2) | 3 | 0.3 |
| Hetero + mutant | 159 | 15.9 |

Abbreviation: *n* Number.

## 4. Discussion

The advances in genomics and high throughput technologies will soon have a profound impact on the management of cardiovascular medicine [44]. Genomics, the science of studying the genes of a genome and how they interact with each other, is at the foundation of personalized medicine, a form of medicine that uses patient’s genomic information to improve diagnosis, prevention, and therapy [45]. Our study was aimed to find a relationship between patients’ genomic profile and their status of polymorphism with reduced scope for aspirin and clopidogrel metabolism. The outcome demonstrated for the presence of 35.9% wild type and 47.2% hetero and 16.9% mutant people who carried CYP2C19*2 variant allele which went in line with the studies conducted in the Malaysian [46], Chinese [47], Egyptian [48] and American [49] healthy population (Table 3).

**Table 3.** Comparison of minor allele frequencies of studied polymorphisms of CYP2C19 gene in different ethnic populations

| Gene variant | Genotype | Egyptian | African | Asian | European | American |
| --- | --- | --- | --- | --- | --- | --- |
| CYP2C19*2 | G>A | 0.126 | 0.169 | 0.334 | 0.148 | 0.115 |
| CYP2C19*3 | G>A | 0.0025 | 0.004 | 0.051 | 0.000 | 0.000 |
| CYP2C19*6 | G>A | 0.000 | - | - | - | - |
| CYP2C19*8 | C<T | 0.000 | 0.004 | 0.003 | - | - |
| CYP2C19*10 | C>T | 0.000 | - | - | - | - |
| CYP2C19*17 | C>T | 0.17 | 0.215 | 0.019 | 0.228 | 0.205 |

Information from 1,000 Genomes [47]. Abbreviation: *CYP* Cytochrome.

While comparing our findings with those from the table 3, it’s kind of shocking to witness a combined percentage of 64.1 in CYP2C19*2 variant allele in our population while most of the CVD patients are taking clopidogrel as an indispensable medicine in their daily medication list. On the other hand, the percentage of CYP2C19*17 carriers (22.7) does not also constitute a very insignificant value. These data symbolized a massive percentage of CYP2C19 poor-metabolizers and rapid-metabolizers in the samples taken.

It was deduced by Sani *et al*. that clopidogrel non-responders were found not only in the carriers of CYP2C19^*^2/^*^2 but also in the carriers of CYP2C19^*^1/^*^2 and CYP2C19^*^1/^*^1 [46]. The potential risk of adverse cardiovascular outcomes among patients with a “poor metabolizer” genotype (for subjects with 2 loss-of-function alleles) advocated for the use of other antiplatelet medications or alternative dosing strategies for them. The investigations of treatment choices associated with more optimal platelet inhibition has led to the inclusion of increasing clopidogrel dosing, adding a third antiplatelet agent (e.g., cilostazol), and switching to a novel generation P2Y12 inhibitor (e.g., prasugrel, ticagrelor, cangrelor) [50]. Hulot *et al*. suggested a doubling of the standard maintenance dose of clopidogrel (75 mg/day) in patients with the CYP2C19*2 loss-of-function polymorphism [52].

In the case of *17 carriers, a standard or reduced dose of clopidogrel and strict monitoring for potential bleeding of the patient was advised [51]. As stated by the current CPIC guidelines, the following changes are to be made:

1. Switching to prasugrel (antiplatelet medication not affected due to CYP2C19 polymorphisms) in poor metabolizer genotype which improves clinical outcome;
2. Discontinuing or lowering clopidogrel doses in rapid metabolizer genotype to decrease bleeding risk [51].

In another exploration in China with 600 patients who underwent PCI, personalized antiplatelet therapy based on CYP2C19 genotype, significantly decreased the incidence of major adverse cardiovascular events (MACE) [52]. However, some studies did not find any momentous decrease in thromboembolic complications in patients whose clopidogrel dosages were tailored as per their demands [49].

The effectiveness of these new drugs (prasugrel, ticagrelor, and elinogrel) as compared to clopidogrel was demonstrated in a handful of studies. These new agents such as prasugrel, ticagrelor, and elinogrel do not undergo CYP2C19 metabolism. Ticagrelor was found to be more effectual for acute coronary syndrome than clopidogrel, irrespective of CYP2C19 and ABCB1 polymorphisms. The researchers comprehended that the use of ticagrelor instead of clopidogrel eliminates the necessity for currently recommended genetic testing before dual antiplatelet treatment [53]. In patients with HTPR after PCI, prasugrel is more effective compared with high clopidogrel in reducing platelet reactivity, particularly in CYP2C19*2 carriers [54].

Chen *et al*.claimed that the effect of low-dose aspirin on CYPs was enzyme-specific after two 7-day and 14-day trials while low-dose aspirin-induced the in vivo activities of CYP2C19 but did not put an impact on the activities of CYP1A2, CYP2D6, and CYP2E1. They also declared that the effect of low-dose aspirin on the CYP3A enzyme anticipates further confirmation. When low-dose aspirin is administered in combination with substrates of CYP2C19, doses of the substrate drugs should also be adjusted to ensure their efficacy [55].

In the case of treatment with aspirin alone, our study did not show a significant plausibility of failure. This result could be backed up by similar studies done to analyze the sensitivity pattern of the same two drugs; aspirin and clopidogrel [56-60]. Lev *et. al*., also found no substantial evidence for an association between these variants and the response towards clopidogrel and aspirin, attributable to the assessment of platelet aggregation and induction markers or by classifying the patients into two different groups based on their drug-sensitivity pattern [61]. As the P2Y12 receptor performed a noteworthy job in platelet aggregation evidenced by a study where this gene was inhibited by a selective antagonist MRS2179 [62], the presence of any variant alleles in this receptor can result in higher induction of platelet activity in cardiovascular patients undergoing PCI.

## 5. Conclusions

The percentages of hetero and mutant variants of the CYP2C19 gene among patients were quite significant, but for the ITGB3gene, it was not that large a number. However, still, our findings from this study surely suggest for patients genotype checking before prescribing any of these drugs. This is applicable for other drugs where a major metabolic gene is responsible for the activation of the drugs as polymorphism can also affect their efficacy to a greater extent unknowingly.

## Abbreviations

ACS: Acute coronary syndrome;
AGE: Agarose gel electrophoresis;
CPIC: Clinical pharmacogenetics implementation consortium;
CVD: Cardiovascular disease;
CYP: Cytochrome;
HTPR: High on-treatment platelet reactivity;
MI: Myocardial infarction;
PCI: Percutaneous coronary intervention.

## Acknowledgment

The authors are thankful to the patients, volunteers, nurses, physicians, and scientists of Labaid Limited (Diagnostic) and Department of Cardiology, United Hospital. The author(s) received no financial support for the research, authorship, and/or publication of this article.

## Authors’ contributions

Study conception and design: TTN, IJ, TMCB, and ZN; Sample collection: NAMM, TTN, and IJ; Data analysis: TTN, IJ, and SNIR; Manuscript writing, editing, and reviewing: TTN, IJ, MSI, RI, AH, and Zn. Supervision of the whole study: RI, AH, and Zn. All authors approved the final version of this manuscript.

## Availability of data and materials

The data used in this study can be available from the corresponding author on a reasonable request.

## Ethics approval and consent to participate

This study was approved by the ethical review committee (ERC), Department of Cardiology, United Hospital, Dhaka, Bangladesh. Before the study, written informed consent was obtained from all subjects included in this study in a prescribed form. The study was conducted according to the Declaration of Helsinki.

## Consent for publication

Not applicable.

